# Dashing: Fast and Accurate Genomic Distances with HyperLogLog

**DOI:** 10.1101/501726

**Authors:** Daniel N Baker, Ben Langmead

## Abstract

Dashing is a fast and accurate software tool for estimating similarities of genomes or sequencing datasets. It uses the HyperLogLog sketch together with cardinality estimation methods that are specialized for set unions and intersections. Dashing summarizes genomes more rapidly than previous MinHash-based methods while providing greater accuracy across a wide range of input sizes and sketch sizes. It can sketch and calculate pairwise distances for over 87K genomes in 6 minutes. Dashing is open source and available at https://github.com/dnbaker/dashing.

## Background

Since the release of the seminal Mash tool [1], data sketches such as MinHash have become instrumental in comparative genomics. They are used to cluster genomes from large databases [1], search for datasets with certain sequence content [2], accelerate the overlapping step in genome assemblers [3, 4], map sequencing reads [5], and find similarity thresholds characterizing species-level distinctions [6]. Whereas MinHash was originally developed to find similar web pages [7], here it is being used to summarize large genomic sequence collections such as reference genomes or sequencing datasets. A collection is reduced to a set of representative *k*-mers and ultimately stored as a list of integers. The summary is much smaller than the original data but can be used to estimate relevant set cardinalities such as the size of the union or the intersection between the *k*-mer contents of two genomes. From these cardinalities one can obtain a Jaccard coefficient (*J*) or a “Mash distance,” which is a proxy for Average Nucleotide Identity (ANI) [1]. These make it possible to cluster sequences and otherwise solve massive genomic nearest-neighbor problems.

MinHash is related to other core methods in bioinformatics. Minimizers, which can be thought of as a special case of MinHash, are widely used in metagenomics classification [8] and alignment and assembly [9]. More generally, MinHash can be seen as a kind of Locality-Sensitive Hashing (LSH), which involves hash functions designed to map similar inputs the same value. LSH has also been used in bioinformatics, including in homology search [10] and metagenomics classification [11].

Spurred by MinHash’s utility, other groups have proposed alternatives using new ideas from search and data mining. BinDash [12] uses a b-bit one-permutation rolling MinHash to achieve greater accuracy and speed compared to Mash at a smaller memory footprint. Other theoretical improvements are proposed in the HyperMinHash [13] and SuperMinHash [14] studies.

Some studies have pointed out shortcomings of MinHash. Koslicki and Zabeti argue that MinHash cardinality estimates suffer when the sets are very different sizes [15]. This is not an uncommon scenario, e.g. when finding the distance between two genomes of very different lengths or when finding the similarity between a short sequence (say, a bacterial genome) and a large collection (say, deep-coverage metagenomics datasets).

Here we use the HyperLogLog (HLL) sketch [16] as an alternative to MinHash that exhibits excellent accuracy and speed across a range of scenarios, including when the input sets are very different sizes and when the sketch data structures are quite small. HLL has been applied in other areas of bioinformatics, e.g. to count the number of distinct *k*-mers in a genome or data collection [17, 18, 19]. We additionally use recent theoretical improvements in cardinality estimation for set unions and intersections [20], the components needed to estimate J and other similarity measures.

We implemented the HLL in the Dashing software tool (https://github.com/dnbaker/dashing), which is free and open source under the GPLv3 license. Dashing supports the functions available in similar tools like Mash [1], BinDash [12] and Sourmash [21]. Dashing can build a sketch of an input sequence set (`dashing sketch`), including FASTA files (for assembled genomes) or FASTQ files (for sequencing datasets). Dashing has a sketch-based facility for removing *k*-mers that likely contain sequencing errors prior to sketching. The `dashing dist` function performs all-pairwise distance comparisons between pairs of datasets in a large collection, e.g. all the complete genomes from the RefSeq database. Since Dashing’s `sketch` function is extremely fast, Dashing can perform both sketching and all-pairs distance calculations in the same command, obviating the need to store sketches on disk between steps. Dashing is parallelized and we show that it scales efficiently to 100 threads. Dashing also uses Single Instruction Multiple Data (SIMD) instructions on modern general-purpose computer processors to exploit the finer-grained parallelism inherent in HLL computations.

## Results

Here we discuss Dashing’s design, then present simulation results demonstrating HLL’s accuracy relative to other data structures. We then describe experiments demonstrating Dashing’s accuracy and computational efficiency relative to Mash and BinDash in a range of scenarios.

Unless otherwise noted, experiments were performed on a Lenovo x3650 M5 system with 4 2.2Ghz Intel E5-2650 CPUs with 12 cores each and 512 GB of DDR4 RAM. Input genomes and sketches were stored on a SAS-attached Lenovo Storage E1000 disk array with 12 8TB 7,200-RPM disks combined using RAID5. All experiments were conducted using scripts available in the `dashing-experiments` repository at https://github.com/langmead-lab/dashing-experiments.

### Design

Dashing uses the HyperLogLog (HLL) sketch to solve genomic distance problems. Dashing takes one or more sequence collections as input. These could be assembled genomes in FASTA format or sequencing datasets in FASTQ format. It then builds an HLL sketch for each input collection based on its *k*-mer content. The sketch can be written to disk or simply forwarded to the next phase, which performs a distance comparison between one or more pairs of sketches. Dashing prints a set of similarity estimates, including estimates for Jaccard coefficient and ANI. It can operate on a given pair of datasets, or can perform all-pairs comparisons across many datasets in a single invocation of the tool.

Dashing is written in C++14. It uses OpenMP for multithreading, with both the sketching and distance phases readily scaling to 100 simultaneous threads. It also uses data-parallel SIMD instructions, including the recent AVX512-BW extensions that have been effective at accelerating other bioinformatics software [22]. Dashing has Python bindings that enable other developers to use the HLL implementation.

### Sketch accuracy

To assess HLL’s accuracy, we measured Jaccard-coefficient estimation error across a range of set and sketch sizes. We implemented both the HLL [16] and MinHash [7] structures in Dashing v0.1.2. For HLL, we used Ertl’s Maximum Likelihood Estimator (MLE) to estimate set cardinalities [20], though we explore alternate methods in later sections. For MinHash, we used a k-bottom sketch with a single hash function, following Mash’s strategy [1]. In both cases we used Thomas Wang’s 64-bit reversible hash function [23]. In both cases, the tools used canonicalized *k*-mers, so that a *k*-mer and its reverse complement are treated as equal when sketching. For details on the commands used to obtain the results, see Additional File 1, Note S1.

We performed several experiments varying (a) the sizes of the two input sets, (b) the degree of overlap between the sets (to achieve a target *J*), and (c) the size of the sketch data structures. Though the structures differ in character, with the HLL storing an array of narrow integers and the MinHash storing an array of wider integers, we can parameterize them to use the same number of bytes of storage. We populated each structure using its natural insert operation; for the HLL this involves hashing the item and using the resulting value to identify the target register and possibly update it according to the leading zero count of the remainder bits (detailed below in Methods). For the bottom-*k* MinHash, inserting involves hashing the item and updating the sketch if the hash is less than the current greatest sketch element. We populated the input sets with random numbers, thereby simulating an ideal hash function with uniformly distributed outputs. Sets were constructed to have target Jaccard coefficients ranging from 0.00022 to 0.818. Many set-size pairs were evaluated ranging from equal-size sets to sets with sizes differing by a factor of 2^12^. In total, we evaluated 36 combinations of set size and *J* were evaluated, with full results presented in Additional File 2. Note that set size and Jaccard-coefficient are dependent; if set *A* has cardinality *c* times greater than set 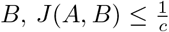.

Figure 1 shows Jaccard-coefficient estimation accuracy results for two values of the true Jaccard coefficient (0.0465 and 0.111) and five pairs of unequal-cardinality input sets. HLL exhibited lower absolute error than MinHash in all cases. We also plotted accuracy for input sets of equal size and for Jaccard coefficients of 0.33, 0.6 and 0.82 (Additional File 1, Figure S1). There, HLL exhibited lower absolute error in most but not all scenarios, with HLL’s greatest advantage coming at smaller sketch sizes.

**Figure 1.**
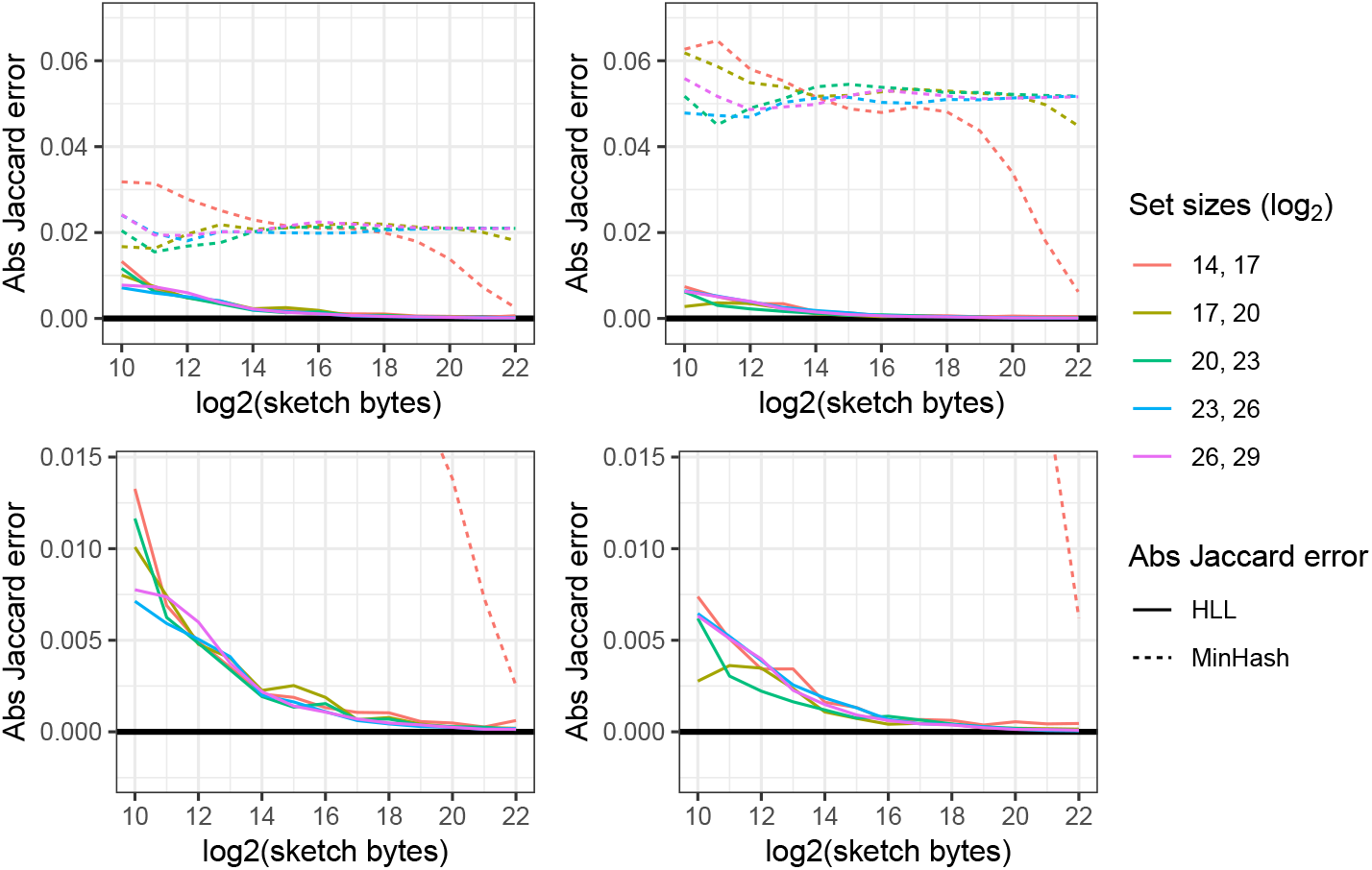
Jaccard-coefficient estimation error using HLL and MinHash. Left column shows experiments with the true Jaccard coefficient fixed at 0.111. Right column shows the same for a coefficient of 0.0465. *x* axis shows the log_2_ of the size of the sketch data structure in bytes. *y* axis shows the absolute error of the Jaccard-coefficient estimate. The second row zooms further in with respect to the *y*-axis. Colors indicate the input set sizes and each pair of inputs differs in size by a factor of 2^3^ = 8.

We also compared HLL and MinHash based sketches to a Bloom-filter based approach [24]. Like HLL and MinHash, a Bloom filter can represent an approximate set, and filters can be compared to estimate union and intersection cardinalities. We implemented and evaluated both a naive (collision-agnostic) and a collision-aware method [25] for estimating set cardinalities via Bloom filters in Dashing v0.1.2. Results are plotted for unequal-cardinality input sets (Additional File 1, Figure S2) and equal-cardinality sets (Additional File 1, Figure S3). Details on our Bloom filter implementation are presented in Additional File 1, Note S2. Bloom-based methods achieved slightly lower absolute error than HLL when the number of bits in the filter approached and exceeded the set cardinality, reflecting the fact that a Bloom-based method eventually converges on error-free “linear counting” given a large enough filter. But HLL exhibited lower error in most circumstances, especially for smaller sketches and larger input sets.

A complete table of results, including all data structures and reporting both absolute and squared errors, can be found in Additional File 2. There we observed that even in adverse scenarios (small data structures and very different set sizes) HLL’s absolute error never exceeded 3% (compared to 8% for MinHash). Overall, the results recommend HLL has an accurate and memory-economical sketch requiring no major assumptions about input set sizes.

### Accuracy for complete genomes

Encouraged by HLL’s accuracy, we measured the accuracy of Dashing v0.1.2’s HLL-based Jaccard-coefficient estimates versus those of Mash v2.1 [1] and BinDash v0.2.1 [12]. We repeated the HLL experiments for three HLL cardinality estimation methods: Flajolet’s canonical method using harmonic mean [16], and two maximum-likelihood-based methods (MLE and JMLE) proposed by Ertl [20]. We selected 400 pairs of bacterial genomes from RefSeq [26] covering a range of Jaccard-coefficient values. To select the pairs, we first used `dashing dist` with *s* = 16, *k* = 31 and the MLE estimation method on the full set of complete bacterial Ref-Seq assemblies (latest versions). We then selected a subset such that we kept 4 distinct genome pairs per Jaccard-coefficient percentile. Our goal was to test an even spread of Jaccard-coefficient values, though some unevenness emerged later due to differences between data structures and different selections of k. Of the genomes included in these pairs, the maximum, minimum and mean lengths were 11.7 Mbp, 308 Kbp, and 4.00 Mbp respectively. For details on exact commands used to obtain and compare the genome pairs, see Additional File 1, Note S3.

We ran the three tools to obtain Jaccard-coefficient estimates for the 400 pairs and plotted the results versus true *J*, as determined using a full hash-table-based *k*-mer counter. Results for *k* = 16 and *k* = 21 and for sketches of size 2^10^ and 2^14^ bytes are shown in Figure 2. The horizontal axis is the true Jaccard-coefficient while the vertical is the difference between the tool-estimated Jaccard-coefficient and the true Jaccard-coefficient . We plot a smooth (loess) curve for each tool to make *J*-dependent biases more evident. For Dashing we used the MLE estimation method. We made a minor change to the Mash software to allow it to output estimated Jaccard coefficient, as it typically emits only Mash distance.

**Figure 2.**
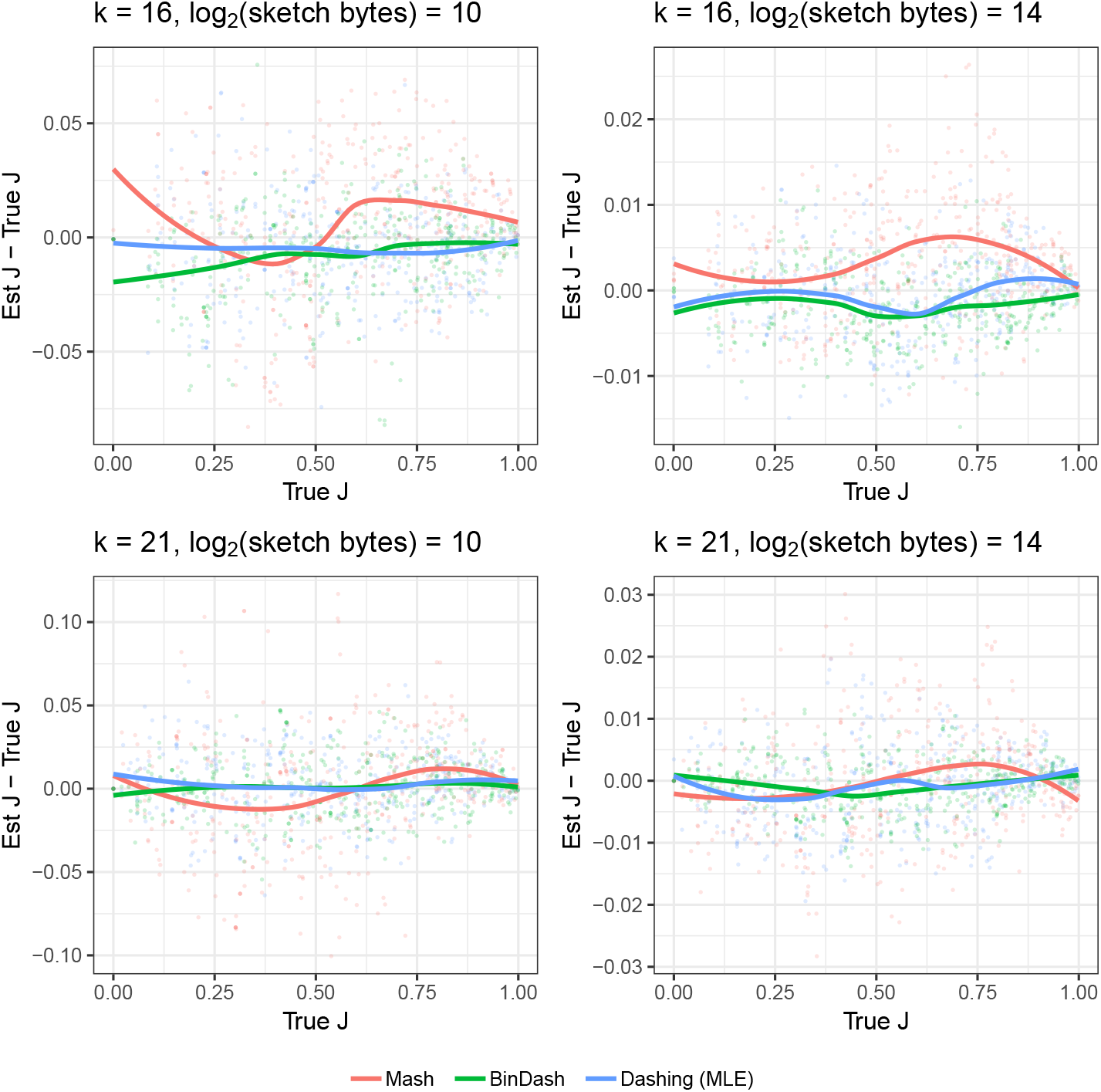
Estimated versus true Jaccard coefficients (*J*s) for various methods across a range of true *J*. Each point is one pair from an overall set of 400 pairs of genomes, selected to evenly cover the range of true *J*s. Smooth loess curves are drawn to highlight systematic bias as a function of *J*.

Dashing’s estimates were consistently near the true *J*. Mash shows a pattern of bias whereby its estimates are somewhat too low at low Jaccard-coefficients then too high at higher coefficients, causing the smooth curve to bend down and then up as it moves to the right. This is sometimes combined with an overall bias shifting estimates too high (in the case of *k* = 16, sketch size = 2^14^) or low (in the case of *k* = 21, sketch size = 2^14^). BinDash and Dashing exhibit less *J*-specific bias.

Table 1 shows mean squared Jaccard-coefficient estimation error (MSE) for a range of sketch sizes and for *k* = 31, also including the two alternate cardinality estimation methods for Dashing (Original and JMLE). The lowest MSEs in each row is shown in bold text. BinDash and Dashing consistently achieve a lower MSE than Mash, with BinDash achieving the lowest error in about half the cases. Dashing seems to achieve lower error relative to BinDash more often for the higher deciles.

**Table 1.** Jaccard-coefficient estimation accuracy for Mash, BinDash and Dashing for *k* = 31 and sketch sizes of 2^*i*^ bytes for *i* ∈ {12,13, 14, 15}. Results are stratified by Jaccard-coefficient decile (first column), and within-decile results are combined by taking the mean squared error across the pairs; the number of such pairs is shown in the *n* column. The lowest error in each row is shown in bold.

| $J$ bin | $k$ | $\log_2(\text{size})$ | Mash | BinDash | HLL<br>Original | HLL<br>Ertl-MLE | HLL<br>Ertl-JMLE | $n$ |
| --- | --- | --- | --- | --- | --- | --- | --- | --- |
| 0 – 0.1 | 31 | 12 | 120.7 | **25.4** | 120.8 | 116.3 | 97.3 | 36 |
|  |  | 13 | 75.9 | **14.5** | 95.6 | 89.6 | 72.7 | 36 |
|  |  | 14 | 27.8 | **3.5** | 42.4 | 41.2 | 31.8 | 36 |
|  |  | 15 | 13.7 | **1.8** | 30.0 | 27.4 | 23.5 | 36 |
| 0.1 – 0.2 | 31 | 12 | 443.5 | **85.0** | 179.8 | 177.7 | 135.2 | 46 |
|  |  | 13 | 228.7 | **28.8** | 92.3 | 88.6 | 76.2 | 46 |
|  |  | 14 | 94.5 | **19.7** | 57.6 | 58.4 | 47.6 | 46 |
|  |  | 15 | 34.9 | **8.2** | 17.1 | 15.8 | 14.0 | 46 |
| 0.2 – 0.3 | 31 | 12 | 478.8 | **57.5** | 118.6 | 113.4 | 91.3 | 38 |
|  |  | 13 | 145.5 | 70.8 | 70.4 | **57.8** | **57.8** | 38 |
|  |  | 14 | 138.4 | **25.0** | 31.6 | 30.0 | 27.7 | 38 |
|  |  | 15 | 79.4 | **9.5** | 21.8 | 19.8 | 20.3 | 38 |
| 0.3 – 0.4 | 31 | 12 | 377.7 | **77.6** | 89.9 | 85.6 | 82.3 | 40 |
|  |  | 13 | 260.5 | 73.8 | 63.9 | 61.1 | **59.5** | 40 |
|  |  | 14 | 115.0 | **30.4** | 39.8 | 41.4 | 36.5 | 40 |
|  |  | 15 | 70.5 | 12.6 | **5.6** | 5.8 | 5.7 | 40 |
| 0.4 – 0.5 | 31 | 12 | 220.4 | 102.4 | 130.4 | 114.4 | **100.0** | 41 |
|  |  | 13 | 99.2 | 72.3 | 39.6 | **36.2** | 39.2 | 41 |
|  |  | 14 | 44.4 | **29.0** | 29.3 | 29.5 | 30.7 | 41 |
|  |  | 15 | 54.4 | **12.2** | 15.3 | 14.8 | 14.1 | 41 |
| 0.5 – 0.6 | 31 | 12 | 379.2 | 138.5 | 131.5 | 132.2 | **124.4** | 39 |
|  |  | 13 | 165.4 | 75.9 | 52.3 | 51.9 | **49.1** | 39 |
|  |  | 14 | 74.6 | 40.1 | 17.3 | 17.6 | **16.0** | 39 |
|  |  | 15 | 87.1 | 16.1 | 13.2 | 12.0 | **11.5** | 39 |
| 0.6 – 0.7 | 31 | 12 | 387.3 | 96.6 | 85.8 | **80.4** | 80.7 | 40 |
|  |  | 13 | 206.0 | **42.3** | 48.5 | 47.9 | 46.0 | 40 |
|  |  | 14 | 63.5 | **25.2** | 37.3 | 36.8 | 34.9 | 40 |
|  |  | 15 | 45.9 | **11.1** | 18.0 | 17.8 | 17.2 | 40 |
| 0.7 – 0.8 | 31 | 12 | 166.3 | 99.4 | 99.9 | 87.9 | **85.5** | 40 |
|  |  | 13 | 121.3 | **41.1** | 67.1 | 63.0 | 61.6 | 40 |
|  |  | 14 | 51.4 | **21.8** | 32.2 | 32.9 | 32.6 | 40 |
|  |  | 15 | 33.1 | **9.4** | 12.2 | 12.4 | 12.6 | 40 |
| 0.8 – 0.9 | 31 | 12 | 222.9 | 87.4 | 69.2 | 65.2 | **64.7** | 39 |
|  |  | 13 | 157.8 | 43.6 | 19.4 | 19.0 | **18.9** | 39 |
|  |  | 14 | 55.0 | 24.6 | 12.0 | **10.4** | **10.4** | 39 |
|  |  | 15 | 50.1 | 9.7 | 6.6 | 5.8 | **5.7** | 39 |
| 0.9 – 1 | 31 | 12 | 70.9 | 35.9 | 19.0 | 14.9 | **14.8** | 41 |
|  |  | 13 | 34.9 | **7.7** | 10.6 | 9.1 | 9.0 | 41 |
|  |  | 14 | 22.1 | 4.5 | 5.1 | **4.3** | **4.3** | 41 |
|  |  | 15 | 9.7 | 5.1 | 1.6 | 1.4 | **1.3** | 41 |

Among the Dashing estimation methods, JMLE consistently achieved the lowest MSE. For computational efficiency reasons (discussed later), Dashing’s default estimation method is the MLE, which had only slightly higher error than JMLE. The table in Additional File 3 shows a fuller set of results for *k* ∈ {16, 21, 31} and sketch sizes of 2^*i*^ bytes for *i* ∈ {10,11,12,13,14,15}.

### Computational efficiency

To assess computational efficiency and scalability in a realistic context, we used Dashing v0.1.2, Mash v2.1 and BinDash v0.2.1 to sketch and perform all-pairs distance calculations for 87,113 complete genome assemblies. We obtained the assemblies from Refseq, filtering to include only assemblies marked “latest” and “Complete genome” and without “contig” in the name. The set included genomes from various taxa, spanning viral, archaeal, bacterial and eukaryotic. Genome lengths varied from 288 bases to 4,502,951,408 bases with mean and median lengths of 9.8Mb and 3.8Mb, respectively. The total number of genome-pair distance calculations required for 87,113 assemblies was over 3.79 billion. We repeated the experiment for a range of sketch sizes and *k*-mer lengths. All experiments were performed on a Lenovo x3850 X6 system with 4 2.0Ghz Intel E7-4830 CPUs, each with 14 processor cores. After hyperthreading, the system supports up to 112 simultaneous hardware threads. The system had 1 TB of DDR4 RAM, and ran CentOS 7.5 Linux, kernel v3.10.0. The system was located at and maintained by the Maryland Advanced Research Computing Center (MARCC).

For Dashing, we used `dashing sketch` for sketching and `dashing dist` for pairwise distance calculations. For Mash, we used `mash sketch` and `mash triangle` for the two stages respectively. For BinDash we used `bindash sketch` and `bindash dist`. We also ran each tool in a way that performed sketching immediately followed by all-pairs distance calculations. For Mash, this involves running its `dist andtriangle` command but specifying the sequence files (rather than their sketches) as input. In the case of `dashing dist`, this combined invocation avoids writing any sketches to disk. Mash provides support for this functionality as well, but we were unable to run it successfully for our large experiment.

All tools were configured to use up to 100 simultaneous threads of execution (Dashing: -p 100, Mash: `-p 100`, BinDash: `--nthreads=100`). Since the system supports a maximum of 112 simultaneous threads, 100 was chosen to achieve high utilization while avoiding excessive contention. We used the GNU `time` utility to measure the average number of CPUs utilized, wall time and peak memory footprint for each tool invocation. For details on the exact commands used, see Additional File 1, Note S4.

For Dashing, we repeated the experiment for each of its three cardinality estimation methods: Flajolet’s canonical method (“Original”), Ertl’s Maximum Likelihood Estimator (“Ertl-MLE”) and Ertl’s joint MLE (“Ertl-JMLE”).

Results for *k* = 21 and *k* = 31 are summarized in Figure 3 and a tabular version of the results for *k* = 31 is shown in Table 2. Full tabular results are shown in Additional File 4.

**Figure 3.**
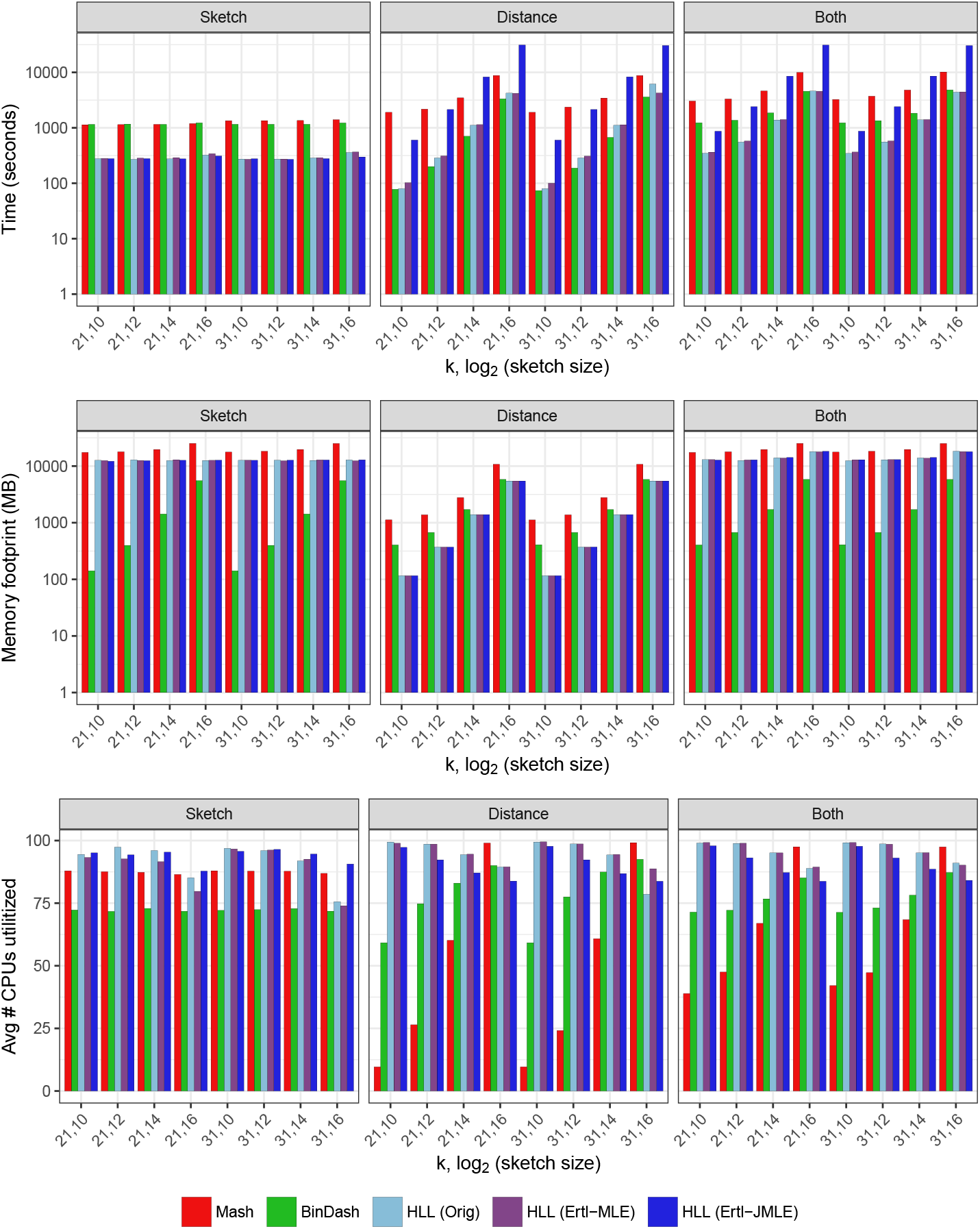
Computational efficiency of Mash, BinDash and Dashing. Results for *k* = 21, *k* = 31 and sketches of size 2^10^ (1KB), 2^12^ (4KB), 2^14^ (16KB) and 2^16^ (64KB). “Both” results obtained either by using a combined Sketch+Distance mode (for Dashing) or by combining results from separate sketching and distance-calculation invocations (for Mash and BinDash). Dashing was assessed using three estimation methods: Flajolet’s method using the harmonic mean (“Orig”) and Ertl’s MLE and JMLE methods.

**Table 2.** Comparison of computational efficiency of Mash, BinDash and Dashing at *k* = 31 and various sketch sizes. The log2(*size*) column reports the log_2_ of the sketch size in bytes. “Both” results obtained either by using a combined Sketch+Distance mode (for Dashing) or by combining results from separate sketching and distance-calculation invocations (for Mash and BinDash). Dashing was assessed using three estimation methods: Flajolet’s method using the harmonic mean (“Original”) and Ertl’s MLE and JMLE methods.

| Phase | Measure | k | $\log_2(\text{size})$ | Mash | BinDash | Dashing<br>Original | Dashing<br>Ertl-MLE | Dashing<br>Ertl-JMLE |
| --- | --- | --- | --- | --- | --- | --- | --- | --- |
| Sketch | Wall clock (s) | 31 | 10 | 1,345 | 1,157 | 273 | **271** | 277 |
|  |  |  | 12 | 1,349 | 1,157 | 273 | 274 | **270** |
|  |  |  | 14 | 1,356 | 1,159 | 286 | 289 | **278** |
|  |  |  | 16 | 1,400 | 1,226 | 359 | 367 | **299** |
|  | Avg # CPUs | 31 | 10 | 87.95 | 72.11 | **96.89** | 96.66 | 95.69 |
|  |  |  | 12 | 87.82 | 72.41 | 95.98 | 96.21 | **96.45** |
|  |  |  | 14 | 87.77 | 72.85 | 91.87 | 92.57 | **94.62** |
|  |  |  | 16 | 86.87 | 71.81 | 75.58 | 73.99 | **90.64** |
|  | Peak mem (MB) | 31 | 10 | 17,720 | **141** | 12,683 | 12,721 | 12,644 |
|  |  |  | 12 | 18,296 | **399** | 12,723 | 12,430 | 12,726 |
|  |  |  | 14 | 19,706 | **1,426** | 12,630 | 12,877 | 12,853 |
|  |  |  | 16 | 25,127 | **5,542** | 12,888 | 12,412 | 12,933 |
| Distance | Wall clock (s) | 31 | 10 | 1,901 | **74** | 80 | 100 | 601 |
|  |  |  | 12 | 2,368 | **188** | 286 | 308 | 2,139 |
|  |  |  | 14 | 3,446 | **672** | 1,113 | 1,137 | 8,308 |
|  |  |  | 16 | 8,777 | **3,603** | 6,172 | 4,251 | 30,506 |
|  | Avg # CPUs | 31 | 10 | 9.66 | 59.19 | 99.4 | **99.48** | 97.72 |
|  |  |  | 12 | 24.15 | 77.53 | **98.64** | **98.64** | 92.33 |
|  |  |  | 14 | 60.82 | 87.43 | 94.28 | **94.45** | 86.83 |
|  |  |  | 16 | **99.17** | 92.51 | 78.57 | 88.76 | 83.76 |
|  | Peak mem (MB) | 31 | 10 | 1,120 | 409 | **116** | **116** | **116** |
|  |  |  | 12 | 1,380 | 673 | **371** | **371** | 372 |
|  |  |  | 14 | 2,785 | 1,709 | **1,392** | **1,392** | **1,392** |
|  |  |  | 16 | 10,776 | 5,816 | **5,476** | **5,476** | **5,476** |
| Both | Wall clock (s) | 31 | 10 | 3,246 | 1,231 | **345** | 365 | 870 |
|  |  |  | 12 | 3,717 | 1,345 | **557** | 579 | 2,407 |
|  |  |  | 14 | 4,801 | 1,831 | **1,390** | 1,408 | 8,574 |
|  |  |  | 16 | 10,177 | 4,829 | **4,394** | 4,453 | 30,433 |
|  | Avg # CPUs | 31 | 10 | 42.09 | 71.34 | 99.17 | **99.22** | 97.72 |
|  |  |  | 12 | 47.26 | 73.13 | **98.64** | 98.56 | 93.04 |
|  |  |  | 14 | 68.43 | 78.2 | 95.1 | **95.17** | 88.59 |
|  |  |  | 16 | **97.48** | 87.26 | 91.01 | 90.18 | 84.12 |
|  | Peak mem (MB) | 31 | 10 | 17,720 | **409** | 12,468 | 12,950 | 12,988 |
|  |  |  | 12 | 18,296 | **673** | 12,958 | 13,042 | 13,020 |
|  |  |  | 14 | 19,706 | **1,709** | 13,951 | 13,782 | 14,205 |
|  |  |  | 16 | 25,127 | **5,816** | 18,320 | 18,081 | 18,011 |

We observed that Dashing is the fastest tool in the Sketch phase, running 3.3-4.3 times faster than BinDash and 3.8-5.0 times faster than Mash. Though Dashing achieves the greatest CPU utilization in the Sketch phase, the tools all perform similarly on this measure, with Mash utilizing about 87 CPUs on average, Bindash about 72 and Dashing about 75-98 (depending on estimation method). Thus, Dashing’s speed is explained mostly by per-thread efficiency.

BinDash achieves the lowest memory footprint among the tools in the Sketch phase, requiring 140 MB for the 1-KB sketch and 5.5 GB for the 64-KB sketch. By contrast, Dashing required about 12 GB across all sketch sizes. This is largely because of how Dashing is parallelized; Dashing threads simultaneously work on separate sequence collections, each filling a buffer of size sufficient to hold the largest sequence yet parsed by that thread. Mash had the highest memory footprint, ranging from 17-25 GB.

In the Distance phase, we noted that the estimation method had a major effect on Dashing’s speed, with JMLE performing 5.9-7.4 times slower than MLE. This is because the JMLE performs significantly more calculations, as described in Methods. This result, together with the relatively small accuracy difference noted earlier, led us to chose the Ertl-MLE method as Dashing’s default. (In a separate experiment, we found that the JMLE inner loop could be made about 20% faster using AVX512BW instructions, as discussed later and detailed in Additional File 1, Note S5 and Table S1.)

BinDash was the fastest tool in the Distance phase, running 25-70% faster than Dashing’s MLE mode. But Dashing is 2-19 times faster than Mash, with the largest speed gap observed at the smallest (1KB) sketch size.

When we compared tools based on combined performance in both the Sketch and Distance phases, BinDash again had the lowest memory footprint (always below 6GB), with Dashing’s footprint at 12-18 GB and Mash’s at 17-25GB. Dashing was the fastest among the three tools at all sketch sizes, though BinDash achieves similar speed at the largest (64KB) sketch size. Mash was the slowest of the tools in all cases. Since small sketch sizes tend to be used in practice (Mash’s default is 4KB or 2^12^ bytes), we expect Dashing to be the fastest overall tool — certainly for sketching, but also combined sketching and distance calculations — in typical situations.

## Discussion

Genomics methods increasingly use MinHash and other locality-sensitive hashing approaches as their computational engines. We showed that the HyperLogLog sketch, combined with recent advances in cardinality estimation, offers a superior combination of efficiency and accuracy compared to MinHash. This is true even for small sketches and for the challenging case where the input sets have very different sizes. While HLL has been used in bioinformatics tools before [17, 18, 19], this is the first application to the problem of estimating genomic distances, the first implementation of the highly accurate MLE and Joint-MLE estimators [20], and the first comprehensive comparison to MinHash and similar methods. The combination of HLL and JMLE is also notable since it directly estimates the cardinality of an intersection, a meaningful quantity independent of its use in the Jaccard coefficient.

We implemented HLL-based sketching and distance calculations in the Dashing software tool. Dashing can sketch and calculate pairwise distances for over 87K Refseq [26] genomes in around 6 minutes using its MLE estimation method, 1KB sketch size, and 100 simultaneous threads of execution (Table 2).

Dashing’s speed advantage is clearest in the sketching step. Notably, re-sketching from scratch is not much slower than loading pre-made sketches from disk. Thus, Dashing users can forgo the typical practice of saving sketches to disk between steps. Dashing’s accuracy with smaller sketches (Figure 1) justifies a lower default sketch size (1KB) compared to Mash’s default of 4KB (or 8KB for long *k*-mers).

It is interesting to observe that Dashing’s accuracy is comparable to that of BinDash across the Jaccard-index deciles in Table 2. Though Dashing is faster — both at sketching and at combined sketching-and-distance — BinDash’s speed approaches that of Dashing at the highest sketch size tested. As we continue to investigate the HyperLogLog sketch, the b-Bit Minwise Hashing technique underlying BinDash is clearly a close competitor, and it will be important to continue to study it as well. In particular, b-Bit Minwise Hashing may also be amenable to SIMD acceleration.

HLL also comes with drawbacks. As shown in Figure 3 and Table 2, Dashing is slower than BinDash at distance calculations. This is expected; the b-bit Minwise Hashing approach consists primarily of comparisons of bit-packed suffixes of minimizers, which can be effectively vectorized. By contrast, the distance calculation between two HLL sketches is relatively expensive, requiring exponentiations, divisions, harmonic means, and — for the MLE-based methods — iterative procedures for finding roots of functions. The trade between accuracy and computational cost is underlined by Ertl’s Joint MLE [20] method, which is both the slowest (even compared to MinHash) but the most accurate of the HLL-based methods. It will be important to continue to refine and accelerate the cardinality-estimation algorithms at the core of `dashing dist`.

HLL lacks another advantage of MinHash; when MinHash is used in conjunction with a reversible hash function, it can be used not only to calculate the relevant set cardinalities but also to report the *k*-mers common between the sets. This can provide crucial hints when the eventual goal is to map a read to (or near) its point of origin with respect to the reference, as is the goal for tools like MashMap [5].

Past efforts have considered how to extend MinHash to include information about multiplicities, essentially allowing it to sketch a multiset rather than a set. This can improve accuracy of genomic distance measurements, especially in the presence of repetitive DNA. Finch [27] works by capturing more sketch items than strictly needed for the k-bottom sketch, then tallying them into a multiset. More theoretical studies have proposed ways to store multiplicities, including BagMinHash [28], and SuperMinHash [14]. In the future it will be important to seek similar multiplicity-preserving extensions — and related extensions like *tf-idf* weighting [3, 29] — for HLL as well.

As we consider how HLL can be extended to improve accuracy and handle multiplicities, an asset is that our current design uses only 6 out of the 8 bits that make up each HLL register. (The LZC of our hash cannot exceed 63 and therefore fits in 6 bits.) Thus, 25% of the structure is waiting for an appropriate use. One idea would be to use the bits to store a kind of striped, auxiliary Bloom filter. This would add an alternate sketch whose strength lies in estimating low-cardinality sets. Since we observed that Bloom filters have superior accuracy when the bitvector is large enough to simulate linear counting (Additional File 1, Figures S2 and S3), we could potentially populate the auxiliary filter with the input items (or a sample thereof) and recover some of the accuracy advantage enjoyed by Bloom filters.

While HLL was used by the KrakenUniq [17] tool for metagenomics read classification, KrakenUniq’s implementation allows for a sparse representation of the registers, with 0-count registers omitted and non-0-count registers stored in a sparse array. Sparsity is a reasonable assumption in KrakenUniq, since some taxa have few associated *k*-mers due to relatedness of the genomes at the leaves. The sparsity assumption is less valid in Dashing’s typical usage scenarios, though it can be valid if one input set has few elements compared to the number of HLL registers. In the future it will be important to investigate whether Dashing can be extended to exploit sparsity where it exists.

Though we compared to Mash and BinDash here, an alternative approach is used by the Kmer-db software [30]. Kmer-db’s data structure captures the *k*-mer content of many input datasets at once. The underlying data structure is a compressed bit matrix with bits indicating membership relationships between *k*-mers (rows) and input datasets (columns). Once a matrix is built, a second phase can perform individual or all-pairwise distance calculations over the samples. Since distinct *k*-mers are represented explicitly — which can take considerable space — the tool gives the option of subsampling the input *k*-mers using a MinHash-based method.

HLL’s accuracy even at low sketch sizes also recommends it as a tool for search and indexing. It can be seen as performing a similar function as the Sequence Bloom Tree [31]. Additionally, because any items which can be hashed can be inserted in a HyperLogLog, dashing could be generalized or extended to other applications, such as comparing text documents by their n-grams, or images by extracted features.

## Methods

### HyperLogLog

The HyperLogLog sketch builds on prior work on approximate counting in 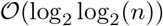 space. Originally proposed by Morris [32] and analyzed by Flajolet [33], this method estimates a count by possibly incrementing a counter with exponentially decaying probability. The probability is typically halved after each increment, so the counter approximates the log_2_ of the true count. While the estimator is unbiased, it has high variance. The hope is that needing only log_2_ log_2_(*n*) bits to store a summary — compared to the log_2_ (*n*) needed for a MinHash — allows us to store more summaries total and, after averaging, achieve a better estimate.

The HLL combines many such counters into one sketch using stochastic averaging [34]. Given a stream of data items, we partition them according to the most significant bits (“prefix”) of their hash values. That is, if o is an input item and h is the hash function, the value *h*(*o*) is partitioned so that *h*(*o*) = *p* ⊕ *q* for bit-string prefix *p* and suffix *q*. To insert the item, we use *p* as an offset into an array of 8-bit “registers.” We update the register to equal either its current value or the leading zero count (LZC) of suffix *q*, whichever is greater (Figure 4a). Note that the LZC of a bit string *x* of length *q* is related to log_2_(*x*):

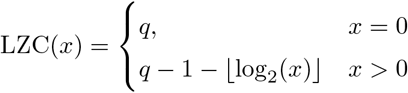

Each register ultimately stores a value related to min_*q*∈*Q*_ log_2_(*q*) where *Q* is the set of suffixes mapping to the register (Figure 4b). We can combine estimates across registers by taking their harmonic mean and applying a correction factor, as detailed below. The estimator has a standard error of 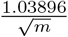.

**Figure 4.**
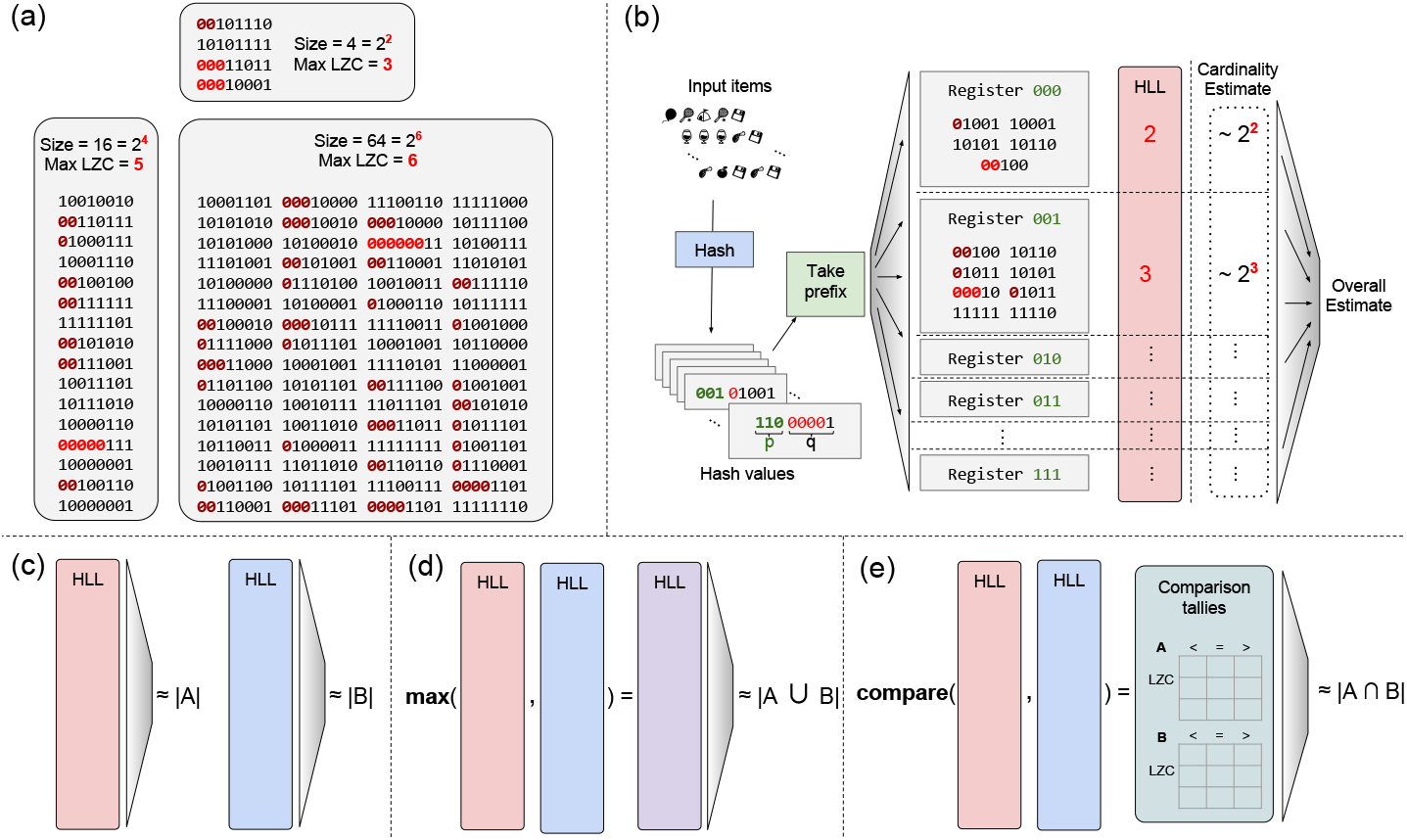
(a) Relationship between maximum leading zero count (Max LZC) and set size for three randomly-generated sets of 8-bit numbers. The Max LZC roughly estimates the log_2_ of the set size, though with high variance; here, two of three estimates are off by 2-fold, (b) Schematic of HyperLogLog sketch. Input items are hashed and hash value is partitioned into prefix *p* and suffix *q. p* indexes into the array of HLL registers. A register contains the maximum leading zero count among all suffixes *q* that mapped there. Register-level estimates are then combined to obtain an overall cardinality estimate, (c) Estimating cardinalities of sets A and B, and (d) estimating the cardinality of their union. For intersection cardinalities using inclusion-exclusion principle, estimated set and union cardinalities are combined, (e) Direct estimation of intersection cardinality with Ert’s JMLE.

While the HLL is conceptually distinct from MinHash sketches and Bloom filters, it is related to both. Informally, an HLL modified so that the summary stored in each register is a simple minimum (without the log_2_) is similar to a MinHash sketch. Similarly, a Bloom filter with a single hash function and 2^*x*^ bits is essentially an HLL with an x-bit hash prefix and with registers consisting of a single bit each.

### Estimation methods

The original HLL cardinality estimation method [16] combines the register-level estimates by taking a corrected harmonic mean:

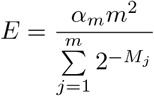

Where *α_m_* is a correction factor equal to 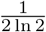 and *M_j_* is 1 + the maximum LZC stored in register *j*. But the estimator’s accuracy suffers at low and high extremes of cardinality. This has spurred various refinements starting with the original HLL publication [16], where linear counting is used to improve estimates for low cardinalities and careful treatment of saturated counters improves high-cardinality estimates.

Ertl proposed further refinements [20]. The “improved estimator” uses the assumptions that (a) the hash function produces uniformly distributed outputs, and (b) register values are independent. It then models register count as a Poisson random variable. Estimating the Poisson parameter yields an estimate for the cardinality.

Ertl’s MLE method again uses the uniformity and Poisson assumptions of the Improved method, but the MLE method proceeds by finding the roots — e.g. using Newton’s method or the secant method — of the derivative of the log-likelihood of the Poisson parameter given the register values. Ertl shows that the estimate is lower- and upper-bounded by harmonic means of the per-register estimates. Ertl suggests using the secant method, which uses inexpensive instructions and avoids derivative calculations. We follow this suggestion in Dashing. Ertl also argues that the MLE generally converges in a small number of steps; we confirm that our implementation converges in at most 3 steps in every case we have tested.

Ertl’s Joint MLE method, unlike those described so far, can directly estimate the cardinality of set intersections. We say “directly” to contrast it with methods that use the inclusion-exclusion principle to estimate intersection cardinality indirectly via cardinalities of sets (Figure 4c) and their unions (Figure 4d). The JMLE method again adopts the Poisson model but two sketches, *A* and *B*, are modeled as a mixture of three components, one with elements unique to A, another with elements unique to B and a third with elements in their intersection *A* ∩ *B*. The method then jointly estimates the Poisson parameters for the three components. The procedure operates on a set of tallies of how often registers having a certain value in *A* are less than, equal to, or greater than their counterparts in *B* (and vice versa) (Figure 4e).

As discussed in Results, the JMLE as implemented in Dashing is substantially slower than MLE. This is partly because of the increased complexity of the numerical optimization, as there are more optimization problems and each requires roughly twice as many iterations as for MLE. However, our profiling indicates the added time is chiefly spent on tallying the <, =, > relationships between the sketch registers. This tallying work grows linearly with the sketch size. This highlights the importance of efficient, SIMD-ized inner loops for comparing HLLs.

We considered but did not include Ertl’s Improved Estimator or the Hyper-LogLog++ estimator [35] in this study as they performed worse than Ertl’s MLE in preliminary comparisons.

### Optimizing speed

Dashing takes advantage of the fine-grained parallelism inherent in HLLs. Union and intersection cardinalities are the key components of similarity measures like the Jaccard coefficient. For two HLLs having the same number of registers and the same hash function, a sketch of their union is simply the element-wise maximum of their registers. Thus, one fundamental need is to perform element-wise maximum over long vectors of 8-bit registers. Finding the cardinatlity of an individual set — or of the intersection of two sets using the JMLE — requires tallying statistics over the register array. Thus, another need is to perform tallies (e.g. counting the registers having a particular value) over long vectors of 8-bit registers.

For set unions, Dashing’s inner loops use Single-Instruction Multiple Data (SIMD) instructions, which are capable of performing fast arithmetic and bitwise operations on vectors of many adjacent operands. These vectors are substantially wider (up to 512 bits) than the typical 32-bit or 64-bit machine words used to store scalar operands. Speedups can be attained by converting important loops to use only or mostly SIMD instructions and to avoid loops with scalar instructions. The more operands per SIMD vector, the greater the potential benefit [22]. The ideal would be to use vectors consisting of 8-bit operands, since this matches the HLL register width. While past iterations of the SIMD instruction set operated on 128- and 256-bit vectors of 8-bit operands, only with the recent introduction of Intel’s AVX-512BW instruction set did it become possible to operate on 512-bit vectors of 8-bit operands. We created AVX-512BW versions of inner set-union loops and confirmed that these deliver the greatest distance-estimation throughput, providing 20% speed boost compared to loops based on the older SSE2 SIMD instruction set (Additional File 1, Note S5 and Table S1). For compatibility with older systems, Dashing supports older SIMD instruction sets back to SSE2.

The process of tallying statistics for set cardinalities and set intersection cardinalities is harder to SIMD-ize in this way. Dashing uses manual loop unrolling to speed up these inner loops, but no SIMD instructions. A question for future work is whether these loops can be rewritten using, for example, a combination of SIMD gather, increment and scatter operations.

Dashing also supports use of many simultaneous threads of execution using the OpenMP v4.5 library. The `dashing sketch` function is parallelized across input files, with distinct threads reading, sketching, and writing sketches for distinct inputs. In `dashing dist`, threads work in parallel on elements in a row of the upper-triangular matrix while a distinct thread writes out the results. To minimize the overhead associated with global memory-allocation locks, each thread allocates from a private memory buffer. The all-pairs distance calculation uses multiple output buffers and asynchronous I/O to avoid blocking and output-lock contention.

Another concern is load balance; having many simultaneous threads is beneficial only if we can avoid “straggler” threads that run long after the others have finished. We eliminated an important source of stragglers by performing an up-front large-to-small ordering of the inputs to be sketched. This minimizes the chance that the thread with the largest genome will still be working when others are finishing.

### Sketching sequencing data

While Dashing supports both FASTA and FASTQ inputs, input data from sequencing experiments require special consideration due to the presence of sequencing errors. Following the strategy of Mash [1], Dashing uses an auxiliary data structure at sketching time to remove infrequent *k*-mers that are likely to contain errors. Dashing does this in a single pass. Each *k*-mer in a sequencing experiment is added to a Count-min Sketch (CMS) [36], and only if the estimated count for the *k*-mer is sufficiently high is it added to the HLL. The CMS can provide count estimates using an amount of space that grows sublinearly with the number of items.

### Hash function

We compared clhash, Murmur3’s finalizer, and the Wang hash across a set of synthetic Jaccard index estimates, and found that Wang’s had the lowest error (8.20 × 10^-3^) and bias (-2.14 × 10^-4^), compared to 8.27 × 10^-3^ and 2.30 × 10^-4^ for Murmur3 and 8.21 × 10^-3^ and —2.66e × 10^-4^ for clhash. In addition to providing the best results, the Wang hash was also much faster than clhash, which is meant for string inputs rather than specialized for 64-bit integers.

## Supporting information

Additional File 2: Full results for sketch Accuracy

Additional File 3: Full results for accuracy for complete genomes

Additional File 4: Full results for computational efficiency

## Competing interests

The authors declare that they have no competing interests.

## Author’s contributions

DNB conceived the method and implemented the software. DNB and BL designed the experiments and wrote the paper. All authors read and approved the manuscript.

## Acknowledgements

We thank Florian Breitwieser for HLL implementation discussions, and Nikita Ivkin for insights with regard to sketch data structure theory and implementation. This work used the Extreme Science and Engineering Discovery Environment (XSEDE), supported by National Science Foundation grant number ACI-1548562.

## Grant support

BL and DNB were supported by National Science Foundation grant IIS-1349906 to BL and National Institutes of Health/National Institute of General Medical Sciences grant R01GM118568 to BL. Experiments on the Intel Skylake system used the XSEDE Stampede 2 resource at the Texas Advanced Computing Center (TACC), accessed using XSEDE allocation TG-CIE170020 to BL.

## Data and software availability

- Dashing source code is available under the open source GPLv3 license at https://github.com/dnbaker/dashing.
- The particular version of Dashing evaluated here is tagged at https://github.eom/dnbaker/dashing/releases/tag/v0.1.2.
- Scripts and code used to perform the experiments described in this study are available under the open source GPLv3 license at https://github.com/langmead-lab/dashing-experiments.
- The particular version of the scripts and code used to perform the experiments described in this study is tagged at: https://github.com/langmead-lab/dashing-experiments/releases/tag/v0.1.
- Accessions of genomes compared in the “Accuracy for complete genomes” subsection of the “Results” section are listed at: https://github.com/langmead-lab/dashing-experiments/blob/master/accuracy/genomes_for_exp.txt.
- Accessions of genomes compared in the “Computational efficiency” subsection of the “Results” section are listed at: https://github.com/langmead-lab/dashing-experiments/blob/master/timing/filenames.txt.

## Additional Files

1. <u>supplement.pdf: Supplementary notes and figures</u> All supplementary notes and figures appear in this additional file.
2. <u>AddTab1.xlsx: Full results for sketch Accuracy</u> Full results from the experimental comparison of MinHash, Bloom, Bloom+, and HyperLogLog methods for Jaccard-coefficient estimation on synthetic data. Results are presented in a spreadsheet.
3. <u>AddTab2.xlsx: Full results for accuracy for complete genomes</u> Jaccard coefficient estimation accuracy across a range of true Jaccard values for BinDash, Mash and 3 HyperLogLog estimation algorithms in tabular format. Experiments were repeated for all combinations of k *E* {16, 21, 31} and log2 sketch size E {10, 11, 12, 13, 14, 15}. Results are presented in a spreadsheet.
4. <u>AddTab3.xlsx: Full results for computational efficiency</u> Space and time efficiency benchmark for all pairwise comparisons between 87,113 genomes for k E {16, 21, 31} and log_2_ sketch size E {10, 14, 16} between BinDash, Mash, and 3 HyperLogLog estimation algorithms. Results are presented in a spreadsheet.

## References

1. Ondov, B.D., Treangen, T.J., Melsted, P., Mallonee, A.B., Bergman, N.H., Koren, S., Phillippy, A.M.: Mash: fast genome and metagenome distance estimation using MinHash. Genome Biol. 17(1), 132 (2016)

2. Schaeffer, L., Pimentel, H., Bray, N., Melsted, P., Pachter, L.: Pseudoalignment for metagenomic read assignment. Bioinformatics 33(14), 2082–2088 (2017)

3. Koren, S., Walenz, B.P., Berlin, K., Miller, J.R., Bergman, N.H., Phillippy, A. M.: Canu: scalable and accurate long-read assembly via adaptive k-mer weighting and repeat separation. Genome Res. 27(5), 722–736 (2017)

4. Berlin, K., Koren, S., Chin, C.S., Drake, J.P., Landolin, J.M., Phillippy, A.M.: Assembling large genomes with single-molecule sequencing and locality-sensitive hashing. Nat. Biotechnol. 33(6), 623–630 (2015)

5. Jain, C., Koren, S., Dilthey, A., Phillippy, A.M., Aluru, S.: A fast adaptive algorithm for computing whole-genome homology maps. Bioinformatics 34(17), 748–756 (2018)

6. Jain, C., Rodriguez-R, L.M., Phillippy, A.M., Konstantinidis, K.T., Aluru, S.: High throughput ANI analysis of 90K prokaryotic genomes reveals clear species boundaries. Nat Commun 9(1), 5114 (2018)

7. Broder, A.Z.: On the resemblance and containment of documents. In: Compression and Complexity of Sequences 1997. Proceedings, pp. 21–29 (1997). IEEE

8. Wood, D.E., Salzberg, S.L.: Kraken: ultrafast metagenomic sequence classification using exact alignments. Genome Biol. 15(3), 46 (2014)

9. Li, H.: Minimap and miniasm: fast mapping and de novo assembly for noisy long sequences. Bioinformatics 32(14), 2103–2110 (2016)

10. Buhler, J.: Efficient large-scale sequence comparison by locality-sensitive hashing. Bioinformatics 17(5), 419–428 (2001)

11. Luo, Y., Yu, Y.W., Zeng, J., Berger, B., Peng, J.: Metagenomic binning through low-density hashing. Bioinformatics (2018)

12. Zhao, X.: Bindash, software for fast genome distance estimation on a typical personal laptop. Bioinformatics, 651 (2018)

13. Yu, Y.W., Weber, G.: Hyperminhash: Jaccard index sketching in loglog space. CoRR abs/1710.08436 (2017). 1710.08436

14. Ertl, O.: Superminhash – A new minwise hashing algorithm for jaccard similarity estimation. CoRR abs/1706.05698 (2017). 1706.05698

15. Koslicki, D., Zabeti, H.: Improving min hash via the containment index with applications to metagenomic analysis. bioRxiv (2017). doi:10.1101/184150

16. Flajolet, P., Fusy, E., Gandouet, O., Meunier, F.: HyperLogLog: the analysis of a near-optimal cardinality estimation algorithm. In: Jacquet, P. (ed.) AofA: Analysis of Algorithms. DMTCS Proceedings, vol. DMTCS Proceedings vol. AH, 2007 Conference on Analysis of Algorithms (AofA 07), pp. 137–156. Discrete Mathematics and Theoretical Computer Science, Juan les Pins, France (2007). https://hal.inria.fr/hal-00406166

17. Breitwieser, F.P., Baker, D.N., Salzberg, S.L.: KrakenUniq: confident and fast metagenomics classification using unique k-mer counts. Genome Biol. 19(1), 198 (2018)

18. Crusoe, M.R., Alameldin, H.F., Awad, S., Boucher, E., Caldwell, A., Cartwright, R., Charbonneau, A., Constantinides, B., Edvenson, G., Fay, S.e.a.: The khmer software package: enabling efficient nucleotide sequence analysis. F1000Res 4, 900 (2015)

19. Georganas, E., Buluc, A., Chapman, J., Oliker, L., Rokhsar, D., Yelick, K.: Parallel de bruijn graph construction and traversal for de novo genome assembly. In: Proceedings of the International Conference for High Performance Computing, Networking, Storage and Analysis. SC ‘14, pp. 437–448. IEEE Press, Piscataway, NJ, USA (2014)

20. Ertl, O.: New cardinality estimation algorithms for hyperloglog sketches. CoRR abs/1702.01284 (2017). 1702.01284

21. Brown, C.T., Irber, L.: sourmash: a library for minhash sketching of dna. The Journal of Open Source Software 1(5) (2016)

22. Rahn, R., Budach, S., Costanza, P., Ehrhardt, M., Hancox, J., Reinert, K.: Generic accelerated sequence alignment in SeqAn using vectorization and multi-threading. Bioinformatics 34(20), 3437–3445 (2018)

23. Wang, T.: Integer Hash Function. http://web.archive.Org/web/20071223173210/http://www.concentric.net/~Ttwang/tech/inthash.htm. [Online; accessed by archive July 2017] (1997)

24. Bloom, B.H.: Space/time trade-offs in hash coding with allowable errors. Communications of the ACM 13(7), 422–426 (1970)

25. Swamidass, S.J., Baldi, P.: Mathematical correction for fingerprint similarity measures to improve chemical retrieval. J Chem Inf Model 47(3), 952–964 (2007)

26. O’Leary, N.A., Wright, M.W., Brister, J.R., Ciufo, S., Haddad, D., McVeigh, R., Rajput, B., Robbertse, B., Smith-White, B., Ako-Adjei, D.e.a.: Reference sequence (RefSeq) database at NCBI: current status, taxonomic expansion, and functional annotation. Nucleic Acids Res. 44(D1), 733–745 (2016)

27. Bovee, R., Greenfield, N.: Finch: a tool adding dynamic abundance filtering to genomic minhashing. Journal of Open Source Software 3(22) (2018)

28. Ertl, O.: Bagminhash – minwise hashing algorithm for weighted sets. CoRR abs/1802.03914 (2018). 1802.03914

29. Chum, O., Philbin, J., Zisserman, A., et al.: Near duplicate image detection: min-hash and tf-idf weighting. In: BMVC, vol. 810, pp. 812–815 (2008)

30. Deorowicz, S., Gudys, A., Dlugosz, M., Kokot, M., Danek, A.: Kmer-db: instant evolutionary distance estimation. Bioinformatics 35(1), 133–136 (2019)

31. Solomon, B., Kingsford, C.: Fast search of thousands of short-read sequencing experiments. Nat. Biotechnol. 34(3), 300–302 (2016)

32. Morris, R.: Counting large numbers of events in small registers. Commun. ACM 21(10), 840–842 (1978)

33. Flajolet, P.: Approximate counting: A detailed analysis. BIT Numerical Mathematics 25(1), 113–134 (1985)

34. Flajolet, P., Martin, G.N.: Probabilistic counting algorithms for data base applications. Journal of computer and system sciences 31(2), 182–209 (1985)

35. Heule, S., Nunkesser, M., Hall, A.: Hyperloglog in practice: Algorithmic engineering of a state of the art cardinality estimation algorithm. In: Proceedings of the 16th International Conference on Extending Database Technology. EDBT ‘13, pp. 683–692. ACM, New York, NY, USA (2013)

36. Cormode, G., Muthukrishnan, S.: An improved data stream summary: the count-min sketch and its applications. Journal of Algorithms 55(1), 58–75 (2005). doi:10.1016/j.jalgor.2003.12.001

