## Additional File 1: Supplementary Information for "Dashing: Fast and Accurate Genomic Distances with HyperLogLog"

### Supplementary Notes

### Note S1 Details on sketch accuracy experiment

The program used to perform the experiments described in this study is named:

- `dsexp/dsexp.cpp`

The R Markdown document used to generate figures and tables is named:

- `dsexp/dsexp.Rmd`

Both are located in the GitHub repository at:

- `https://github.com/langmead-lab/dashing-experiments`

Note that this experiment uses Dashing's HLL implementation, as well as Dashing's own implementations of the MinHash sketch and the Bloom filter.

### Note S2 Details on Bloom filter sketches

A Bloom filter [1] can be used to hold an approximate representation of a set. Further, Bloom filters for different sets can be compared to arrive at cardinality estimates for set unions and intersections. To this end, we implemented a Bloom-filter method for estimating Jaccard coefficient in Dashing v0.1.2, in addition to the HLL and MinHash methods evaluated in the main text. We conducted two additional series of simulation experiments to test both a naive (collision-agnostic) and a collision-aware method [2] for estimating set cardinalities from the Bloom filters.

While Bloom filters can use several hash functions to improve accuracy in some situations, we used one hash function in our experiments. This is a reasonable choice in the absence of any foreknowledge of the set cardinality, as it yields the lowest false positive rate for the case where the number of bits in the filter is not much larger than the set cardinality.

Results are shown in Figures S2 and S3. Figure S2 shows results for unequal-sized input sets. For the two $J$s tested (0.0465 and 0.111), HLL had lower absolute error than MinHash as seen most clearly in the second row. The Bloom-filter-based methods, especially the collision aware method (Bloom+), achieved slightly lower error in some scenarios. Bloom+ had lower error (by $< 0.0025$) for sketches of size $2^{13}$ bytes and higher for the smallest sets. For larger sets, the point where Bloom+ began to have lower error than HLL moved rightward. The naive Bloom method also sometimes outperformed HLL, though only for the largest sketch size and smallest input sets. That the Bloom filters outperform at large sketch sizes is not surprising; as the number of filter bits increases far past the set cardinality, collisions become rare and the method converges on error-free linear counting.

Figure S3 shows results for input sets of equal size and for Jaccard coefficients of 0.33, 0.6 and 0.82. HLL again had lower absolute error in most scenarios, though the Bloom-based approaches — similarly to the unequal-input-set case — had slightly lower absolute error for sufficiently large sketches. MinHash also achieved lower error in a few scenarios.

### Note S3 Details on accuracy for complete genomes experiment

The script used to select the 400 genome pairs used is named:

- `accuracy/pairselector.py`

The program used to perform the experiments is named:

- `accuracy/pairwise_benchmark.cpp`

The scripts used to generate figures and tables are named:

- `accuracy/ji_range.Rmd`
- `accuracy/ji_range_postprocess.py`

All of the above are located in the GitHub repository at:

- `https://github.com/langmead-lab/dashing-experiments`

### Note S3.1 Dashing

Rather than calling Dashing by command-line, instead the encoder and sketch structure used in dashing were called directly in C++ as below:

```
enc.for_each([&](u64 v) {
    kh_put(all, hash_, v, &khr);
    v = sketch::hll::hll_t::HashType()(v);
    for(auto &h: hll_) h.add(v);
}, fp);
```

- The same hash and other parameters in Dashing were used.
- The only difference is that each key was only hashed once and then inserted into multiple HLLs to reduce experimental cost.

### Note S3.2 Mash

```
mash sketch -s %zu -k %u -o %s %s
```

```
mash dist -d 0. -j %s %s\n
```

Where:

- `-s %zu` allows `%zu` integers in the final MinHash sketch
- `-k %u` sets the $k$-mer length to `%u`
- `-o %s` directs output to a file named `%s`
- `-d 0.` causes Mash to emit all values, rather than only those passing a filter
- `-j %s` is a special parameter supported by our fork of the Mash project (at `https://github.com/dnbaker/Mash`). It causes Mash to emit Jaccard coefficients rather than Mash distances.

### Note S3.3 BinDash

```
bindash sketch --minhashtype=2 --bbits=16
    --sketchsize64=%zu --kmerlen=%u --outfname=%s %s
```

```
bindash dist --mthres=1e300 %s %s
```

Where:

- `--minhashtype=2` uses the (default) method of $b$-bit MinHash with optimal densification.
- `--bbits=16` sets the parameter $b$ for $b$-bit MinHash to be 16.
- `--sketchsize64=%zu` requests `%zu` 64-bit integers to be used in the sketch.
- `--outfname=%s` directs output to `%s`.
- `--kmerlen=%u` sets the $k$-mer length to be `%u`.
- `--mthres=1e300` sets the maximum distance to report to be 1e300, effectively requiring that all distances be emitted.

### Note S4 Details on computational efficiency experiment

The script used to perform the experiments described in this study is named:

- `timing/all_pairwise.py`

The R Markdown document used to generate figures and tables is named:

- `timing/timing.Rmd`

And both are located in the GitHub repository at:

- `https://github.com/langmead-lab/dashing-experiments`

#### Note S4.1 Mash

The command we used to perform sketching experiments was:

```
nohup {time_path} -v mash sketch -p {threads}
    -s {1<<(sketchsz - 2 - (ksz > 16))}
    -k {ksz} -o mash.{ksz}.{sketchsz}.msh
    -l {fname_paths} &> {mashlog}
```

- Where `{nohup}` is a utility that ensures the command is not interrupted even if the user subsequently logs out
- `{time_path}` is the path to a utility which is used for full timing information. (GNU time 1.9) `-v` emits verbose information.
- `{threads}` is the number of simultaneous hardware threads to use
- `{kzs}` is the $k$-mer length
- `{sketchsz}` is the $\log_2$ of the sketch size in bytes
- The expression `{1<<(sketchsz - 2 - (ksz > 16))}` selects number of minimizers to request from Mash to result in a sketch of a given power of 2 size in bytes. The `ksz > 16` clause accounts for Mash's use of 32-bit minimizers when `ksz <= 16`.
- Parameter -k sets the k-mer size to be :param:.
- `{fname_paths}`, specified by parameter `-l`, provides the a list of paths, one per line, of genomes to process.

The command we used to perform distance experiments was:

```
nohup {time_path} -v mash triangle -p {threads}
     mash.{ksz}.{sketchsz}.msh
     1> {mashout} 2>> {mashlog}
```

- Where `{nohup}` is a utility that ensures the command is not interrupted even if the user subsequently logs out.
- `{time_path}` is the path to a utility which is used for full timing information. (GNU time 1.9) `-v` emits verbose information.
- `{threads}` is the number of simultaneous hardware threads to use.
- `{mashout}` is the path to write results are written.
- `{mashlog}` is the path to which logging information, including results from $time\_path$, are appended.

### Note S4.2 Dashing

The command we used to perform sketching experiments was:

```
nohup {time_path} -v dashing sketch
{sketchstr} -k{ksz} -p{threads}
-S{sketchsz} -F{fname_paths} &> {logfile}
```

- Where {nohup} is a utility that ensures the command is not interrupted even if the user subsequently logs out.
- {time_path} is the path to a utility which is used for full timing information. (GNU time 1.9) -v emits verbose information.
- {threads} is the number of simultaneous hardware threads to use.
- {sketchstr} is the command-line flag corresponding to which mode to use. (-I for Ertl Improved, -E for original Flajolet, -J for Ertl JMLE, and an empty string for Ertl MLE).
- {kzs} is the $k$-mer length.
- {sketchsz} is the $\log_2$ of the sketch size in bytes.
- {fname_paths}, specified by parameter -F, provides the a list of paths, one per line, of genomes to process.

The command we used to perform distance calculations was:

```
nohup {time_path} -v dashing dist
{sketchstr} -Wfbk{ksz} -p{threads}
-S{sketchsz} -o{outsizes}
-O{outdists} -F{fname_paths} &> {logfile}.tmp
```

- Where {nohup} is a utility that ensures the command is not interrupted even if the user subsequently logs out
- {time_path} is the path to a utility which is used for full timing information. (GNU time 1.9) -v emits verbose information.
- {threads} is the number of simultaneous hardware threads to use
- {kzs} is the $k$-mer length

- {sketchsz} is the $\log_2$ of the sketch size in bytes
- {fname_paths}, specified by parameter -F, provides the a list of paths, one per line, of genomes to process.
- Parameter -O specifies the file to which to write the distance matrix.
- Parameter -o specifies the file to which to write a list of genomes which their estimated cardinalities.
- Parameter -W specifies to use pre-sketched sketches (instead of sketching from the genomes).
- Parameter -f specifies to emit single-precision floats.
- Parameter -b specifies to emit binary format.

The command we used to perform combined sketching and distance calculations was:

```
nohup {time_path} -v dashing dist {sketchstr}
-fbk{ksz} -p{threads} -S{sketchsz}
-o{outsizes} -O{outdists}
-F{fname_paths} &> {logfile}.tmp
```

- Where {nohup} is a utility that ensures the command is not interrupted even if the user subsequently logs out.
- {time_path} is the path to a utility which is used for full timing information. (GNU time 1.9) -v emits verbose information.
- {threads} is the number of simultaneous hardware threads to use
- {kzs} is the $k$-mer length
- {sketchsz} is the $\log_2$ of the sketch size in bytes
- {fname_paths}, specified by parameter -F, provides the a list of paths, one per line, of genomes to process.
- Parameter -O specifies the file to which to write the distance matrix.
- Parameter -o specifies the file to which to write a list of genomes which their estimated cardinalities.
- Parameter -f specifies to emit single-precision floats.

- Parameter `-b` specifies to emit binary format.

Note: the lack of parameter `-W` causes Dashing to perform both sketching and comparisons instead of using cached sketches.

#### Note S4.3 BinDash

The command we used to perform sketching experiments was:

```
nohup {time_path} -v bindash sketch --minhashtype=2
--nthreads={threads} --bbits={BBITS}
--sketchsize64={ns64} --kmerlen={ksz}
--outfname=bindash.{ksz}.{sketchsz}.bdsh
--listfname={fname_paths} &> {bindashlog}
```

- Where `{nohup}` is a utility that ensures the command is not interrupted even if the user subsequently logs out.
- `{time_path}` is the path to a utility which is used for full timing information. (GNU time 1.9) `-v` emits verbose information.
- `{threads}` is the number of simultaneous hardware threads to use.
- `{kzs}` is the $k$-mer length
- `{ns64}` is the parameter designating the number of sets of 64 minimizers to use to match the sketch size of dashing in bytes.
- `{fname_paths}`, specified by parameter `--listfname=`, provides the a list of paths, one per line, of genomes to process.
- `--bbits={BBITS}` sets the parameter $b$ for $b$-bit MinHash. For all our experiments, $b$ was set to 16 to make sketch sizes equal to powers of two possible.
- `{sketchsz}` is the $\log_2$ of the sketch size in bytes.

The command we used to perform distance calculations was:

```
nohup {time_path} -v bindash dist
--nthreads={threads} bindash.{ksz}.{sketchsz}.bdsh
1> {bindashout} 2>> {bindashlog}
```

- Where `{nohup}` is a utility that ensures the command is not interrupted even if the user subsequently logs out

- `{time_path}` is the path to a utility which is used for full timing information. (GNU time 1.9) `-v` emits verbose information.
- `{threads}` is the number of simultaneous hardware threads to use
- `{kzs}` is the $k$-mer length
- `{sketchsz}` is the $\log_2$ of the sketch size in bytes.

### Note S5 SIMD instruction set comparison

We compiled a separate binary for each of four successive iterations of the Intel SIMD instruction set. The first binary used only instructions up to and including the SSE2 instruction set, introduced in the 2000 Pentium 4 architecture and supporting vector words of 128 bits. The second binary used instructions up to and including the AVX2 instruction set, introduced in the 2013 Haswell architecture and supporting 256-bit words. The third used instructions up to and including the AVX512 instruction set, introduced in the 2016 Knight's Landing architecture and supporting 512-bit words (with a caveat described below). The fourth used instructions up to the AVX512BW instruction set, which was expanded to include instructions with 8- and 16-bit operands.

To experiment with all of these instruction sets on the same computer, we used a single Skylake node from the Stampede 2 supercomputer at the Texas Advanced Compute Center (TACC). The specific node had dual-socket Intel Xeon Platinum 8160 processors, each with 24 cores and 2 hyperthreaded threads per core, for a total of 96 simultaneous hardware threads. Each core was clocked at a 2.1 GHz nominal rate and the node had 192GB of DDR4 RAM.

The difference between AVX512 and AVX512BW is that AVX512 lacks instructions for vectors of 8-bit operands; AVX512 instructions work on vectors or 32- or 64-bit operands only. For Dashing this means we cannot use 512-bit instructions that map naturally onto the 8-bit HyperLogLog registers and instead must use (AVX2) instructions operating on 256-bit-wide vectors of registers. We still use 512-bit instructions to load and store portions of the HLL and thereby derive some benefit from the 512-bit vector words.

Table S1 shows CPU-time and wall-clock time results for the distance calculation step only, and for the four instruction sets and for Ertl's Maximum Likelihood Estimator (MLE) and Joint MLE (MLE). The distance calculation was the same all-pairs calculation between 87,113 complete genome assemblies as was benchmarked in the "Computational efficiency" Results section in the main text. The exact Dashing command used was the as for the "Computational efficiency" experiment (see Note S4), but using 90 hardware threads (`-p90`) instead of the 100 threads we used in other experiments, since the Skylake node supports only up to 96 simultaneous threads. Here we measure the results for a sketch size of 1KB and a $k$-mer length of 31 (`-S10 -k31`).

For JMLE, the wider vector words generally led to faster distance calculations, with AVX512BW performing best overall and about 20% faster than SSE2. The AVX512 method, which used 512-bit loads and stores but otherwise relied on AVX2 instructions, performed worse than both our AVX512BW version and our pure AVX2 version. For MLE, wider vector words led to slower distance calculations, with SSE2 performing best overall. This is likely due to the fact that Intel's frequency scaling technology reduces the processor clock frequency when wider vector words are used [3]. We hypothesize that the decrease in clock speed leads to a net slowdown for MLE (versus a net speedup for JMLE) because the MLE inner loop has a lower density of vector arithmetic compared to JMLE.

Given that the MLE estimation method is Dashing's default, the released Dashing binaries — and the default setting when a user compiles the Dashing binary from scratch — use the SSE2 instruction set by default. Given the efficacy of AVX512BW for the JMLE method, it will be important to investigate how to include both SSE2 and AVX512BW modes in a single binary executable without impairing portability.

### Supplementary Tables

| Method | Instruction set | Vector width | CPU time (thread-seconds) | Wall time (seconds) |
|---|---|---|---|---|
| JMLE | SSE2 | 128 | 45,288 | 515 |
| | AVX2 | 256 | 37,420 | 431 |
| | AVX512 | 512 | 39,109 | 456 |
| | AVX512BW | 512 | **35,818** | **410** |
| MLE | SSE2 | 128 | **8,151** | **105** |
| | AVX2 | 256 | 8,819 | 106 |
| | AVX512 | 512 | 9,725 | 123 |
| | AVX512BW | 512 | 9,268 | 118 |

Table S1: Running time of Dashing all-pairs distance calculation between 87,113 complete genome assemblies for various SIMD instruction sets.

### Supplementary Figures

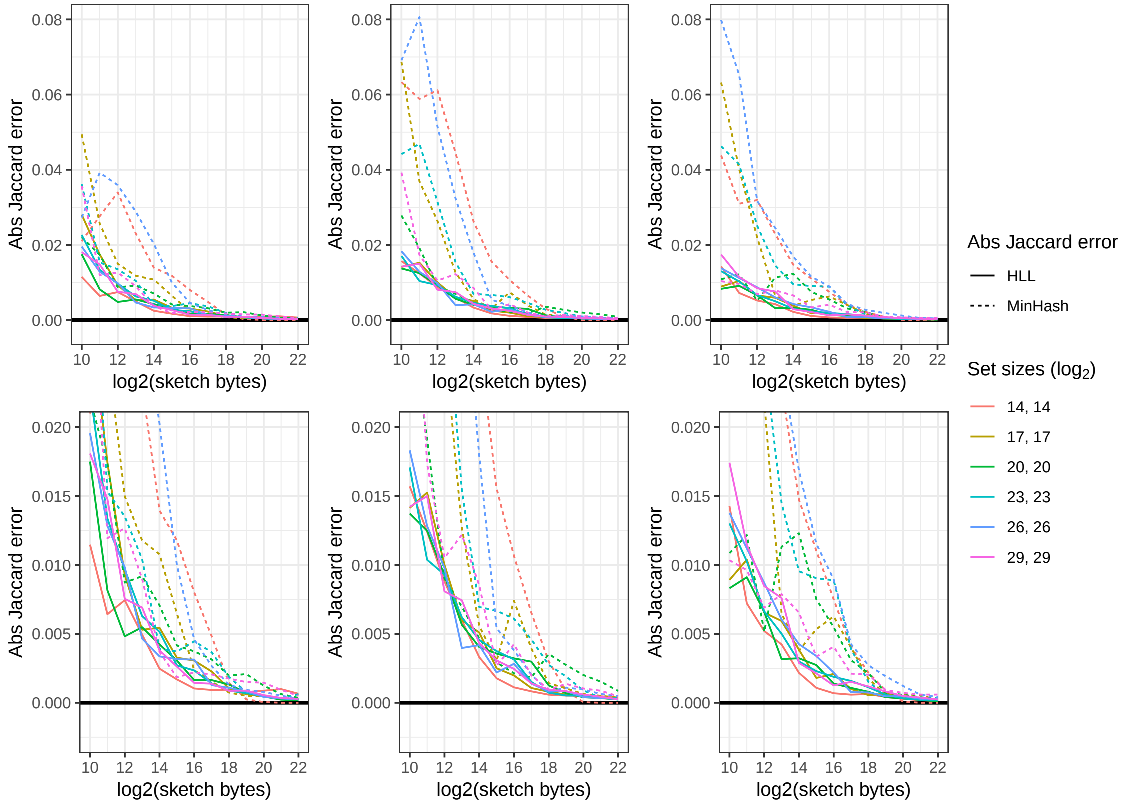


Fig S1: Columns correspond to simulations where the true Jaccard coefficients were 0.33, 0.6 and 0.82 (respectively, left to right). The $x$ axis shows the $\log_2$ of the sketch size in bytes. The $y$ axis shows the absolute error of the method. All input-set pairs are equal-size, with size labeled by color. The second row zooms further in with respect to the $y$-axis.

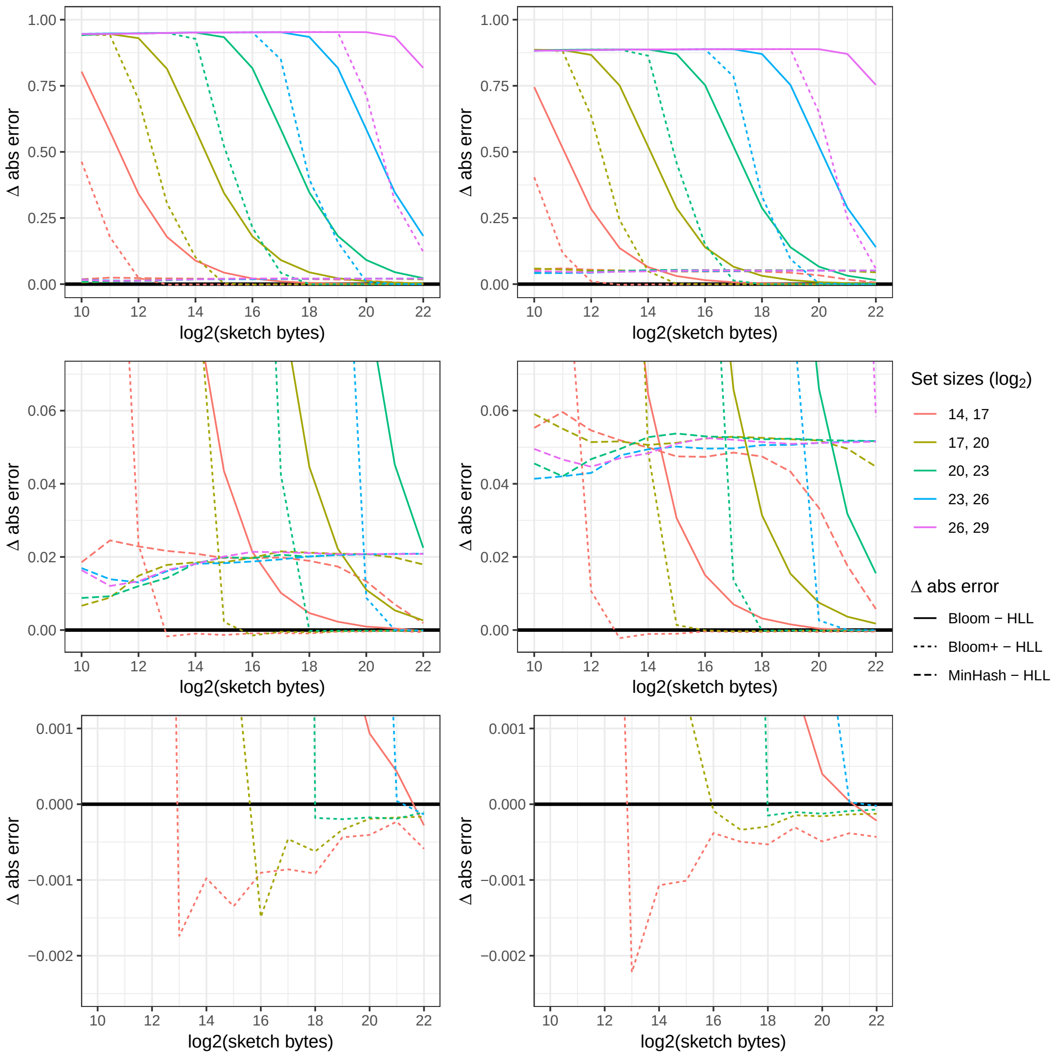


Fig S2: Columns correspond to simulations where the true Jaccard coefficients were 0.111 (left) and 0.0465 (right). The $x$ axis shows the $\log_2$ of the sketch size in bytes. The $y$ axis shows the absolute error of the alternative method minus that of the HLL; points above the $y$ axis indicate HLL's error is lower than that of the alternative method. All pairs of input sets differ in size by a factor of $2^3 = 8$. The second and third rows zoom further in with respect to the $y$-axis so as to highlight the relative performance of MinHash (2nd row) and the Bloom+ filter (3rd row). HLL exhibits lower absolute error in most circumstances, especially for smaller sketches and larger sets. Bloom+ starts to outperform ($\Delta$ absolute error drops below $y = 0$) when the number of bits in the filter is sufficiently large compared to the cardinality. "Bloom+" refers to the collision-aware estimation method, whereas "Bloom" refers to the naive method.

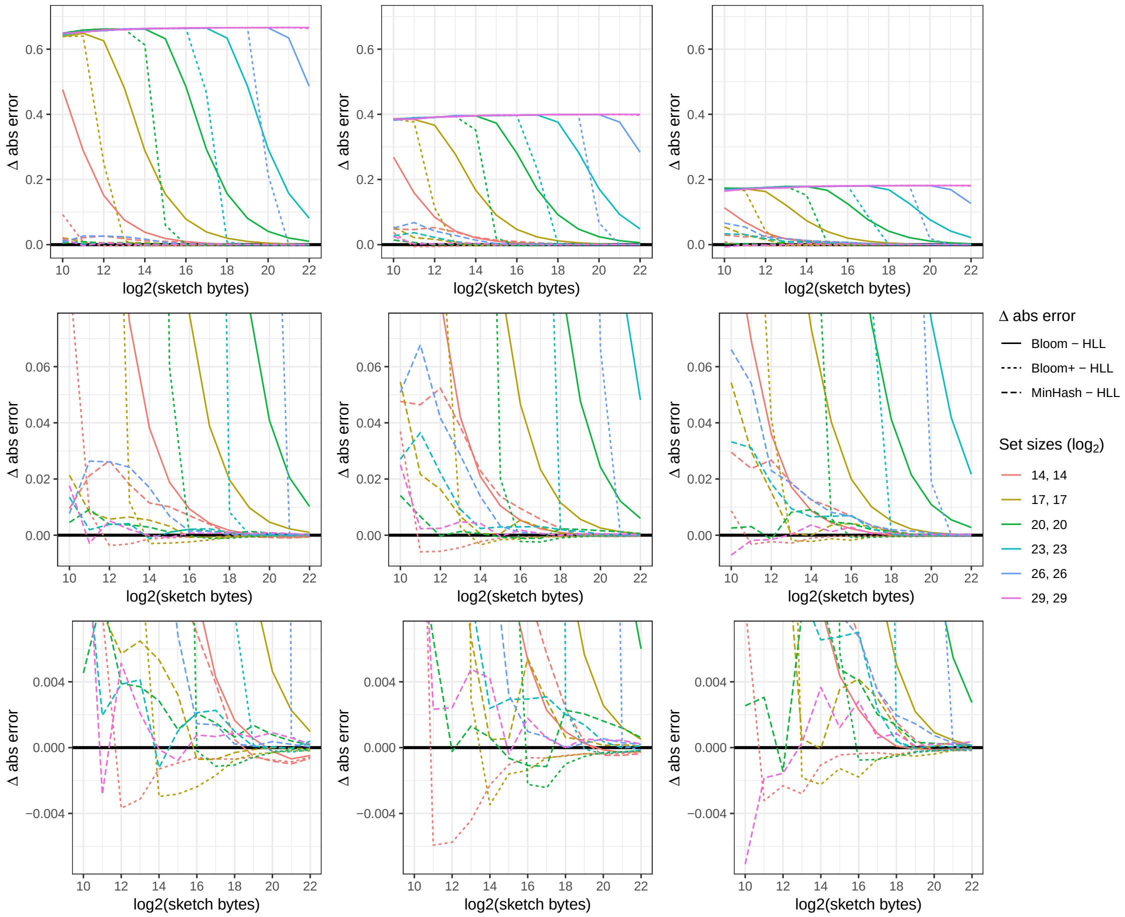


Fig S3: Columns correspond to simulations where the true Jaccard coefficients were 0.33, 0.6 and 0.82 (respectively, left to right). The $x$ axis shows the $\log_2$ of the sketch size in bytes. The $y$ axis shows the absolute error of the alternative method minus that of the HLL; points above the $y$ axis indicate HLL's error is lower than that of the alternative method. All pairs of input sets are the same size, with size labeled by color. The second and third rows zoom further in with respect to the $y$-axis. "Bloom+" refers to the collision-aware estimation method, whereas "Bloom" refers to the naive method.

### References

[1] Burton H Bloom. "Space/time trade-offs in hash coding with allowable errors". In: *Communications of the ACM* 13.7 (1970), pp. 422–426.

[2] S. J. Swamidass and P. Baldi. "Mathematical correction for fingerprint similarity measures to improve chemical retrieval". In: *J Chem Inf Model* 47.3 (2007), pp. 952–964.

[3] Mathias Gottschlag and Frank Bellosa. "Mechanism to Mitigate AVX-Induced Frequency Reduction". In: *CoRR* abs/1901.04982 (2018). arXiv: `1702.01284`. URL: `https://arxiv.org/abs/1901.04982`.
